# Simulation of cancer cell line pharmacogenomics data to optimise experimental design and analysis strategy

**DOI:** 10.1101/174862

**Authors:** Hitesh Mistry, Phil Chapman

## Abstract

**Background:** Explaining the variability in drug sensitivity across a panel of cell lines using genomic information is a key aspect of cancer drug discovery. The results of such analyses may ultimately determine which patients are likely to benefit from a new treatment. There are numerous experimental factors that can influence the outcomes of cell line screening panels such as the number of replicates, number of doses explored etc. Simulation studies can aid in understanding how variability in these experimental factors can affect the statistical power of a given analysis method. In this study dose response data was simulated for a variety of experimental designs and the ability of different methods to retrieve the original simulation parameters was compared. The analysis methods under consideration were a combination of non-linear least squares and ANOVA, conventional approach, versus non-linear mixed effects.

**Results:** Across the simulation studies explored the mixed-effects approach gave similar and in some situations greater statistical power than the conventional approach. In particular the mixed-effects approach gave significantly greater power when there was less information per dose response curve, and when more cell lines screened. More generally the best way to improve statistical power was to screen more cell lines.

**Conclusions:** This study demonstrates the value of simulating data to understand design and analysis choices in the context of cancer drug sensitivity screening. By illustrating the performance of different methods in different situations these results will help researchers in the field generate and analyse data on future preclinical compounds. Ultimately this will benefit patients by ensuring that biomarkers of drug sensitivity have an increased chance of being identified at the preclinical stage.

## Introduction

Testing anti-cancer compounds on model systems with different genetic backgrounds to assess the correlation of genetic features to compound response is a central tenet of biomarker discovery. The exemplar of this approach is the discovery that loss of *BRCA1* or *BRCA2* conferred an increase in sensitivity to PARP (poly (ADP-ribose) polymerase) inhibitors (1,2) Since then, numerous large scale screens of hundreds of compounds in panels containing up to 1000 cell lines have been conducted. These screening studies are designed to enable hypothesis free discovery of novel biomarkers of drug sensitivity (3–6). Drug discovery groups now routinely screen novel compounds in such panels to generate hypotheses on which patient subgroups are most likely to benefit from the new compound (7,8). Consistency and reproducibility of these projects has been a source of debate (9–15). Efforts have also been made to leverage these datasets for a variety of purposes (16) and provide infrastructure for analysis (17).

Within drug discovery, dose response screening on a large scale predominantly involves testing a variety of compounds within the same biological system. As a result, experimental noise remains approximately the same across compounds. With cell line screening, however, data is being compared from different assays where growth characteristics of cell lines vary. One analysis approach that takes this confounding factor into consideration is to calculate the concentration at which cell growth rate is reduced by 50%, GR50, (18). Other confounding factors that are routinely being accounted for are tissue specific effects (19), and the general level of drug sensitivity (20).

Dose response data is conventionally analysed by carrying out a non-linear regression on each unique combination of compound and assay to generate an estimate of IC50 or Area Under Curve for use in subsequent analyses. This approach discards uncertainty in the estimates, and doesn’t allow information to be shared between curves. The use of mixed effects models where data is combined across cell lines and drugs has been shown to increase the accuracy of IC50 estimates for large scale screens by sharing information (21). An extension of this approach is to include the genetic covariate itself in the non-linear mixed effects model. Estimating the genetic effect in one step rather than two theoretically allows uncertainty information to be retained which may improve precision and reduce bias.

Screens can be designed in different ways: number of cell lines screened, concentration range, number of concentrations tested, and number of replicates per concentration can all be varied. A large scale screen of 1000 cell lines may have a single replicate per concentration and only 8 or 9 different concentrations (3,4) whereas a pharmaceutical company may screen a smaller panel of 20 cell lines in triplicate with 10 different concentrations. Inherent or unknown variables that will affect the ability of the screen to detect genetic effects include the effect size itself, the proportion of cell lines with a feature, the variability in response to compound, the experimental noise, and the efficacy of the compound relative to minimum and maximum dose.

In the present study, cell lines and their dose response data were simulated to compare the ability of different analysis methods and experimental designs to recapture known genetic effects. The results presented will assist researchers in choosing appropriate analysis methods and assist in experimental design strategies to get the best balance of cost and power for the scientific question being asked.

## Methods

### Simulating cell lines

The pIC50 (pIC50 = log_10_ (IC50)) values for wild-type cell lines were sampled from a normal distribution with mean *m*_*1*_ and standard deviation *s*_*1*_. The pIC50 values of the mutant cell lines were sampled from a normal distribution with mean *m*_*1*_-*k*_*1*_ and standard deviation *s*_*1*_. That is only the population mean of the pIC50 values was assumed to change between mutant and wild-type cell lines whereas the variation was assumed to be the same.

### Simulating dose response curves

Combining pIC50 values for the wild-type and mutant cell lines gave us a population of pIC50 values. For each pIC50 value we then simulated a response value, *R*, for a cell line *i (i=1,…,n)* at dose *j (j = 1,…,m)* for replicate *k (k = 1,…,l)* using,

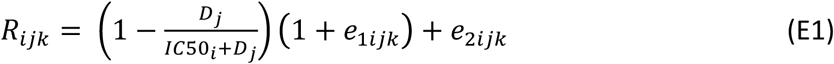

In the above equation *D*_*j*_ refers to the drug concentration value *j* used, *IC50*_*i*_ refers to cell line *i’*s IC50 value, while *e*_*1ijk*_ and *e*_*2ijk*_ are the proportional and additive residual error values sampled from a normal distribution with standard deviation *s*_*2*_ and *s*_*3*_ respectively. These residual error terms perturb the true dose-response to create noisy data. We also created a genotype vector *G*_*i*_, which represents what genotype *IC50*_*i*_ came from, E1 equates to the mutant cell-line and 0 the wild-type.

### Methods for retrieving genetic effect

Given that the data generation process began with sampling pIC50 values the first method we used, which can be considered as a benchmark, was to perform an ANOVA with sampled pIC50 values with genotype as a covariate. We collected the p-value, from the F-test, and the estimated size of the effect together with 95 percent confidence intervals. The R function lm was used for this analysis. Therefore we shall refer to this approach as lm. In the subsequent methods we used the noisy dose-response data generated.

The first of these involved fitting the following dose-response model (E2) to each cell line using the nls.lm function from the minpack.lm R package.

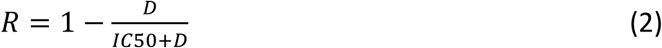

This led to the generation of a distribution of estimated IC50 values which was transformed to generate a distribution of pIC50 values for the population of cell lines. Any pIC50 estimates that fell either below the minimum concentration tested minus 3 log10 units or above the maximum concentration plus 3 log10 units were set to these limits. These estimated pIC50 values were then used within an ANOVA in the same way as the lm approach detailed above. This approach is referred to as nls_lm.

The next method involved fitting the dose-response model described in (E2) within a hierarchical modelling framework (21) using the nlme R package. That is we replace the parameter IC50 in (E2) with,

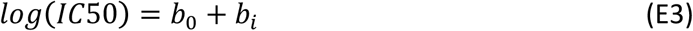

where *b*_*0*_ is the population estimate of the *log(IC50)* value and *b*_*i*_ is the distance from *b*_*0*_ for each cell line *i*. The individual pIC50 values derived from the estimation were then subsequently used within an ANOVA as stated above. This approach is referred to as nlme_lm.

The final method involved modifying equation (E3) to include genotype in the following way,

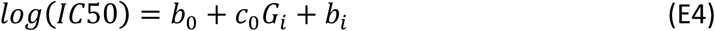

where *c*_*0*_ is the shift in the population estimate of *log(IC50)* for the mutant cell lines versus the wild-type. Model (E3) can be considered to be nested within model (E4) therefore we used the likelihood ratio test to assess if the fit to the data improved with model (E4) over (E3). We collected the p-value and the estimate of *c*_*0*_ and the 95 percent confidence intervals. This approach is referred to as nlme_gene.

### Simulations carried out

Four simulations were carried out to explore different types of experimental design (Table 1).

**Table 1:** Set of simulation parameters used within the studies.

| Sim | mu_pIC50 | sd_pIC50 | Effect size | Prop. | N | sd_add | sd_prop | Doses |
| --- | --- | --- | --- | --- | --- | --- | --- | --- |
| 1 | -5 to 4 | N/A | N/A | N/A | 181 | 0.1, 0.2, 0.4, 0.8 | 0.1, 0.2, 0.4, 0.8 | 7x1, 10x3 |
| 2 | 0,1,2,3 | 0.4, 1 | 1 | 0.5 | 50 | 0.15, 0.5 | 0.3 | 7x1, 10x3 |
| 3 | 0.5 | 0.4, 1 | 0.3, 0.5, 1 | 0.5 | 10, 20, 40 | 0.15, 0.5, 1 | 0.3 | 7x1, 10x3 |
| 4 | 0.5 | 1 | 0.3, 0.5, 1 | 0.05, 0.1, 0.2 | 50, 200, 800 | 0.15, 0.5 | 0.3 | 7x1, 10x3 |

mu\_pIC<sub>50</sub> - population mean pIC<sub>50</sub> value; sd\_pIC<sub>50</sub> - standard deviation of the distribution of pIC<sub>50</sub> values; Prop. – proportion of cell lines that have a mutation; sd\_add – standard deviation of the additive error; sd\_prop – standard deviation of the proportional error

The first simulation compared the actual pIC50 with that estimated by the conventional non-linear regression model (2) and the hierarchical model (3) across a range of pIC50 values and with different amounts of additive and proportional noise.

The second simulation went on to examine how each of the four methods performed at retrieving the genetic effect when the pIC50 population mean was altered but the proportion of mutant cell lines and the number of cell lines was kept the same.

The third simulation kept the proportion mutated at 0.5 and average pIC50 fixed at 0.5 (~3*μ*M), varied numbers of cell lines (10, 20, 40), effect size (0.3, 0.5, 1), additive error (0.15, 0.4, 1), and variation in pIC50 values (0.4, 1).

The fourth simulation was meant to represent a realistic cell line panel screen of different sizes (50, 200, 800 cell lines) with two different amounts of additive error (0.15 and 0.4), three different effect sizes (0.3, 0.5, 1) and three different proportions (0.05, 0.1, 0.2).

For each simulation, two different experimental designs were considered: in the first a 10 point dose response curve with 3 replicates at each dose was simulated (10pt_3rep), whereas in the second a 7 point dose response curve with a single replicate was simulated (7pt_1rep). The former represents ‘gold standard’ data that might be generated at low throughput, whereas the latter represents high throughput data produced in a large scale screen.

For each of the simulation set-ups, 2 to 4, described above and summarised in Table 1 we conducted 200 simulations. The distribution of the results were then explored both graphically and quantitatively via reporting the statistical power; proportion of simulations which gave a p-value <0.05.

### Running the simulations

Simulations were carried out in R version 3.4.0 using the *pgxsim* package (GitHub repo <u>https://github.com/chapmandu2/pgxsim</u>). R scripts for the different simulations and instructions for their use can be found on GitHub at <u>https://github.com/chapmandu2/pgx_simulation_scripts</u>). Amazon Web Service c4.x8large instances were provisioned using the RStudio Amazon Machine Image maintained by Louis Aslett (http://www.louisaslett.com/RStudio_AMI/) and analysis was parallelised using the batchtools R package (Lang et al 2017).

## Results

### Simulation 1: Compare real vs estimated pIC50

10 point triplicate dose response curves simulated with varying amounts of additive and proportional error with an actual pIC50 of 0 (1uM) are shown in Figure 1. Consultation with a number of scientists and showing them these dose response curves gave rise to the consensus view that additive residual error values (sd_add) between 0.4 and 0.8 and proportional residual error value (sd_prop) between 0.2 and 0.4 represented the maximum noise that would be acceptable in a panel screen.

**Figure 1:**
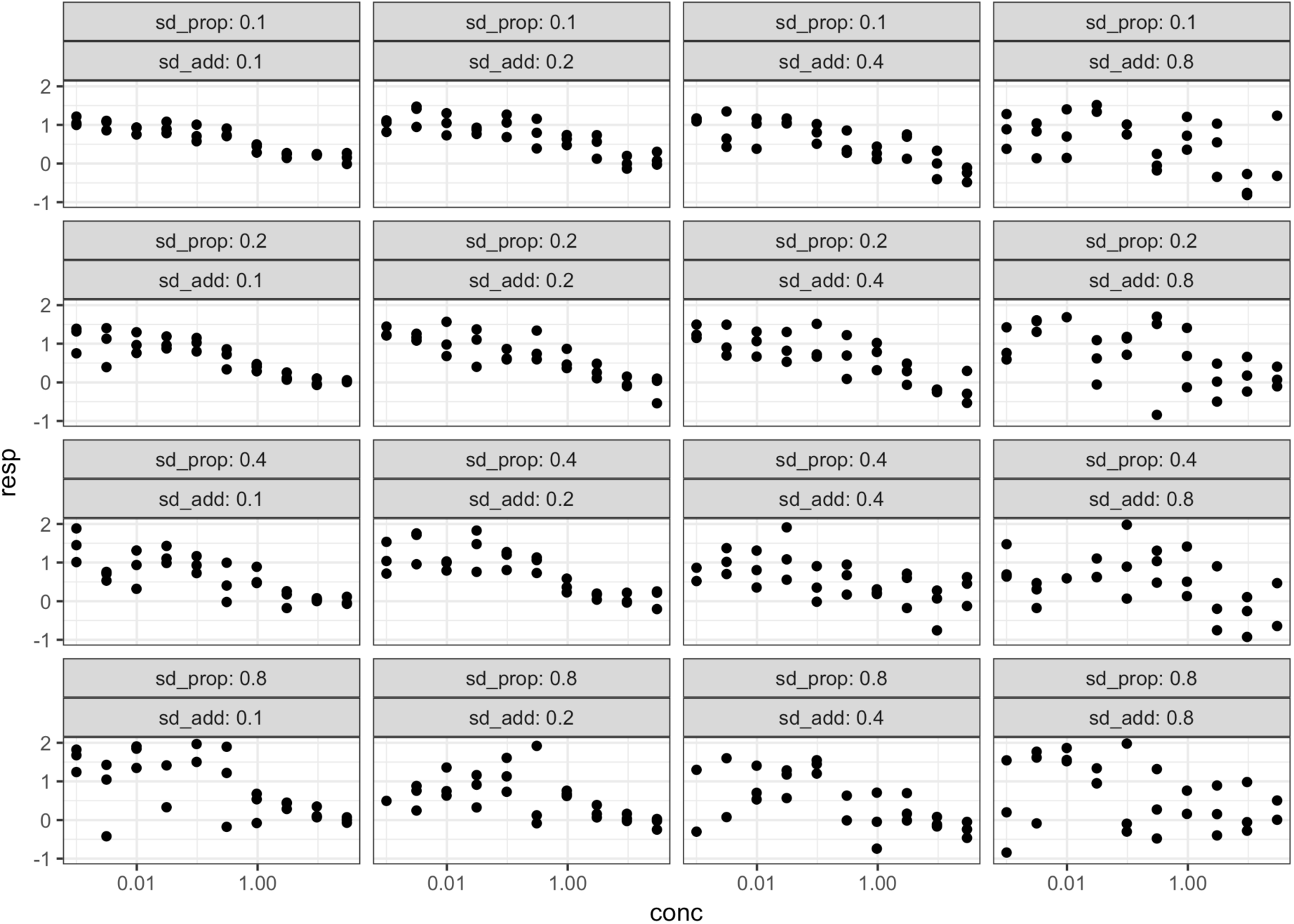
Simulated 10-point triplicate dose response curves from Simulation 1 for a pIC50 value of 0 with different amounts of proportional (sd_prop) and additive (sd_add) variance.

Correlation plots of real versus estimated pIC50 using the nls (non-linear least squares) and nlme (non-linear mixed-effects) methods from 10 point triplicate and 7 point single replicate dose response curves are shown in Figure 2. As expected, the relationship between actual and estimated pIC50 is stronger for the 10 point triplicate than 7 point singe replicate curve, and is also stronger when there is less variability in the simulated data.

**Figure 2:**
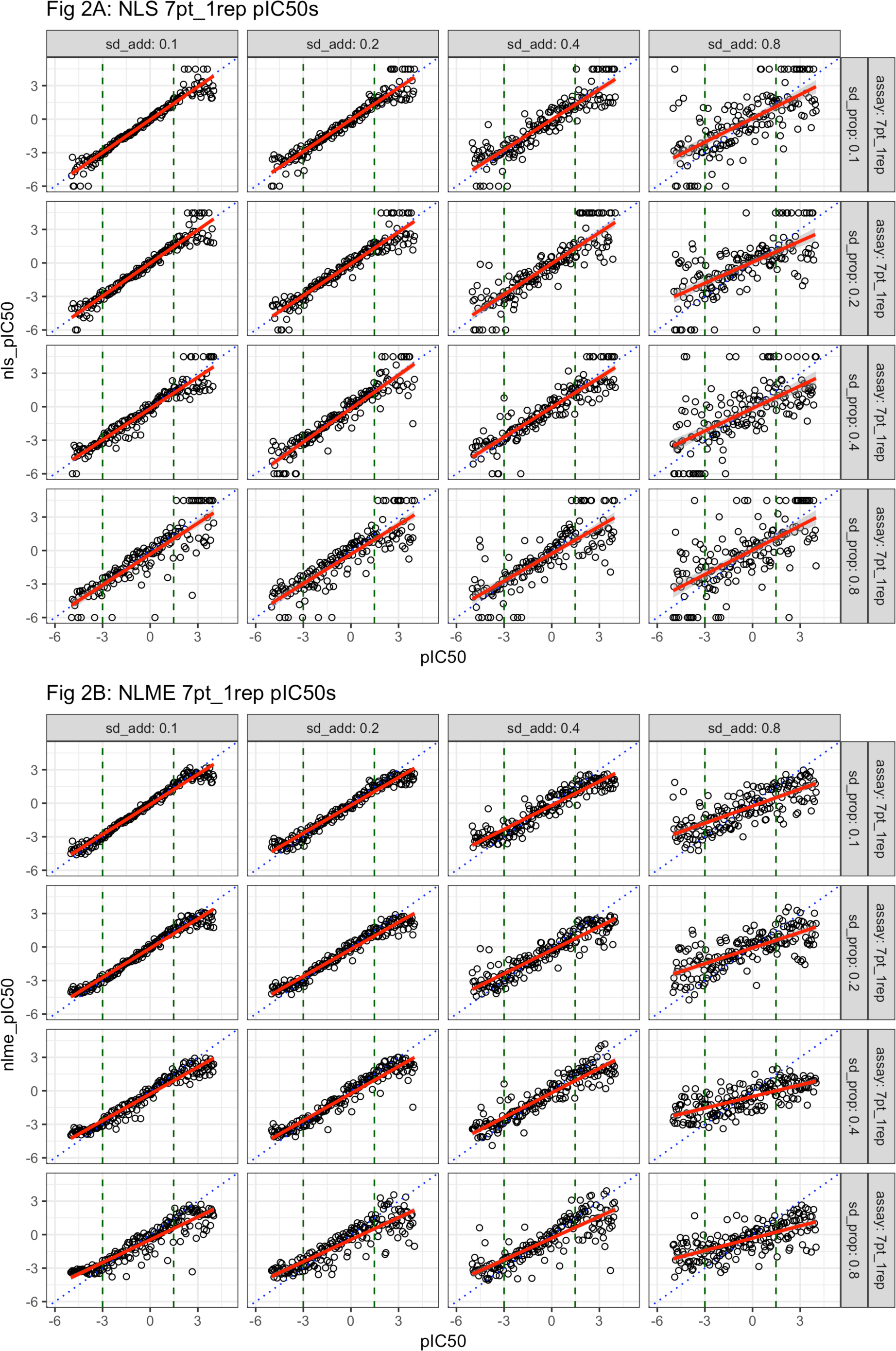
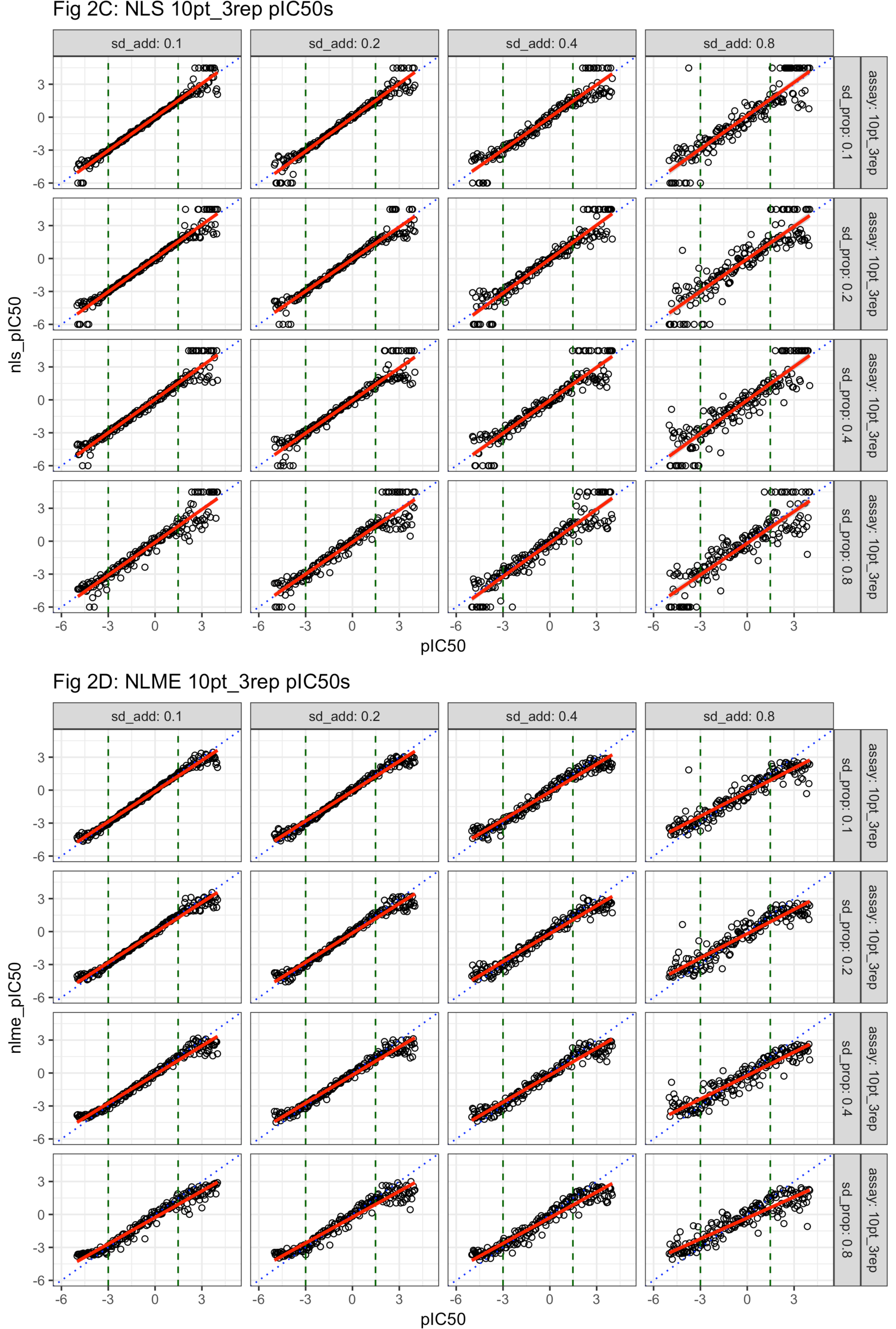
Correlation plot of actual vs estimated pIC50 values from Simulation 1 applying the nls (2A/C) and nlme (2B/D) methods across a range of values of additive (sd_add) and proportional (sd_prop) variance for two types of dose response curve, 7 point single repliicate (2A/B) and 10 point triplicate (2C/D. Blue dotted line is identity line, red solid line is linear fit, green dashed vertical lines represent minimum and maximum concentration of the dose response curve.

Whilst the plots for each model are similar, the most visible effect is how estimates of pIC50’s outside of the experimental dose response range (vertical green broken lines) are handled. The standard model can produce very large or very small estimates which are truncated to a maximum of 3 log units above/below the minimum/maximum concentration. By contrast, when there is less information the mixed effects model regularised the estimates towards the population mean pIC50.

### Simulation 2: Explore effect of mu on estimating genetic effect

Estimated genetic effect sizes and p-values for each simulation are plotted in Figure 3. There is a systematic shrinkage of the estimate of the genetic effect size towards zero for the nlme_lm method, while nlme_gene seems to give slightly more precise estimates than nls_lm. However, the distribution of p-values is similar across all methods, with the nls_lm method showing slightly worse performance under certain circumstances. In particular, all methods perform less well as the average pIC50 moves above and beyond the maximum dose and the difference between the nlme based methods and nls_lm gets more pronounced. There is also a greater difference between methods when there is more variability, and/or more points in the dose response curve.

**Figure 3:**
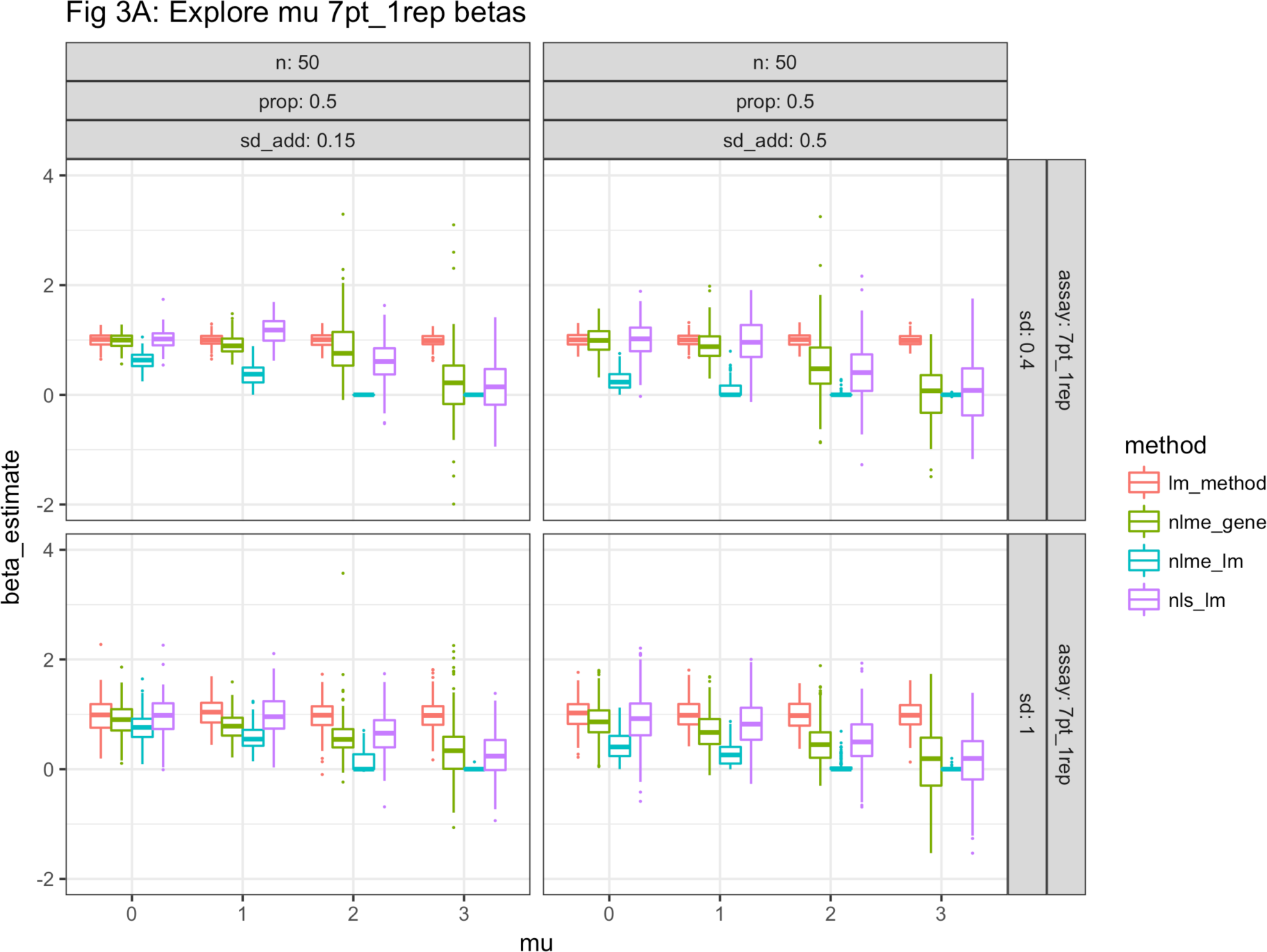
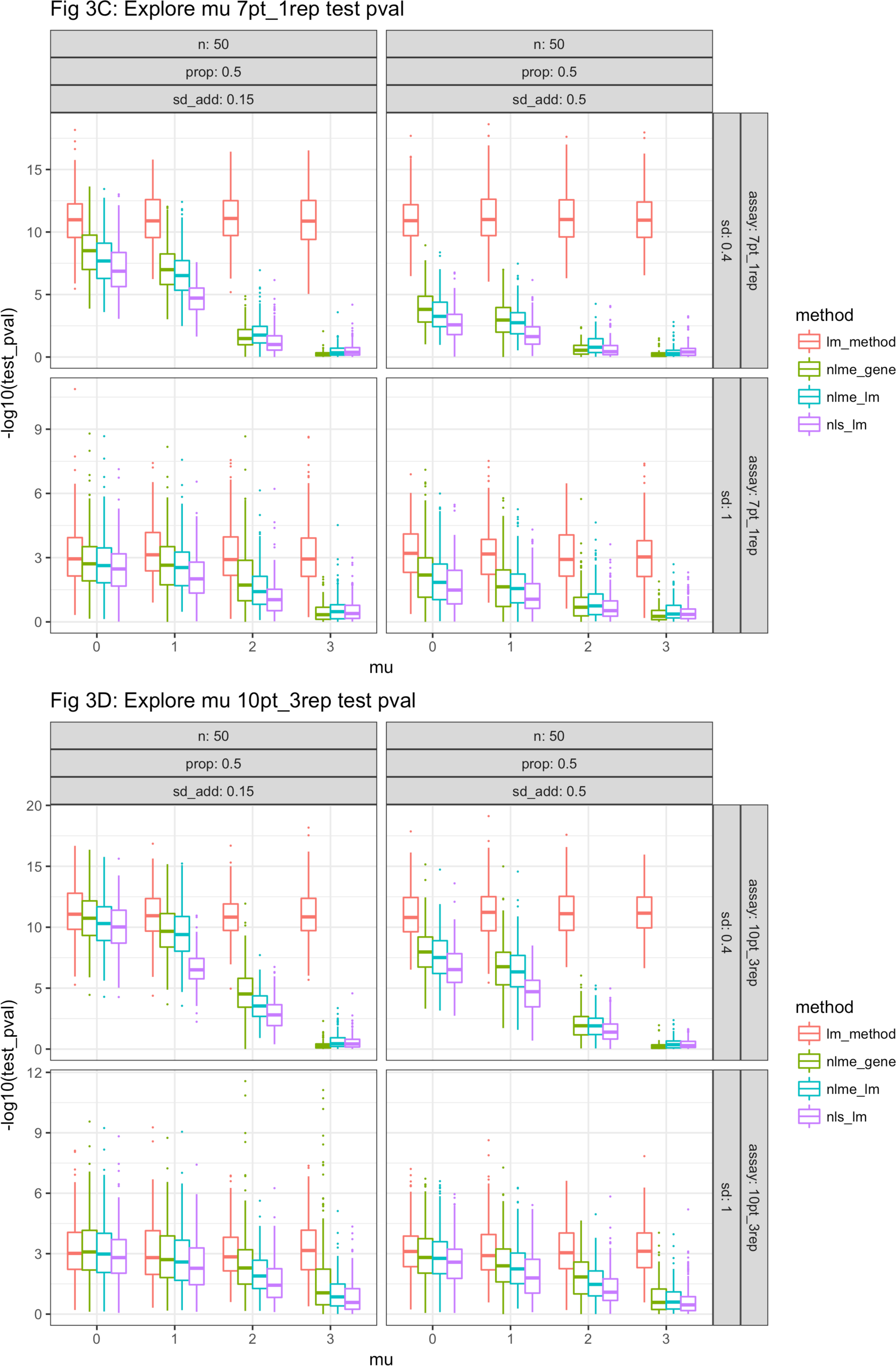
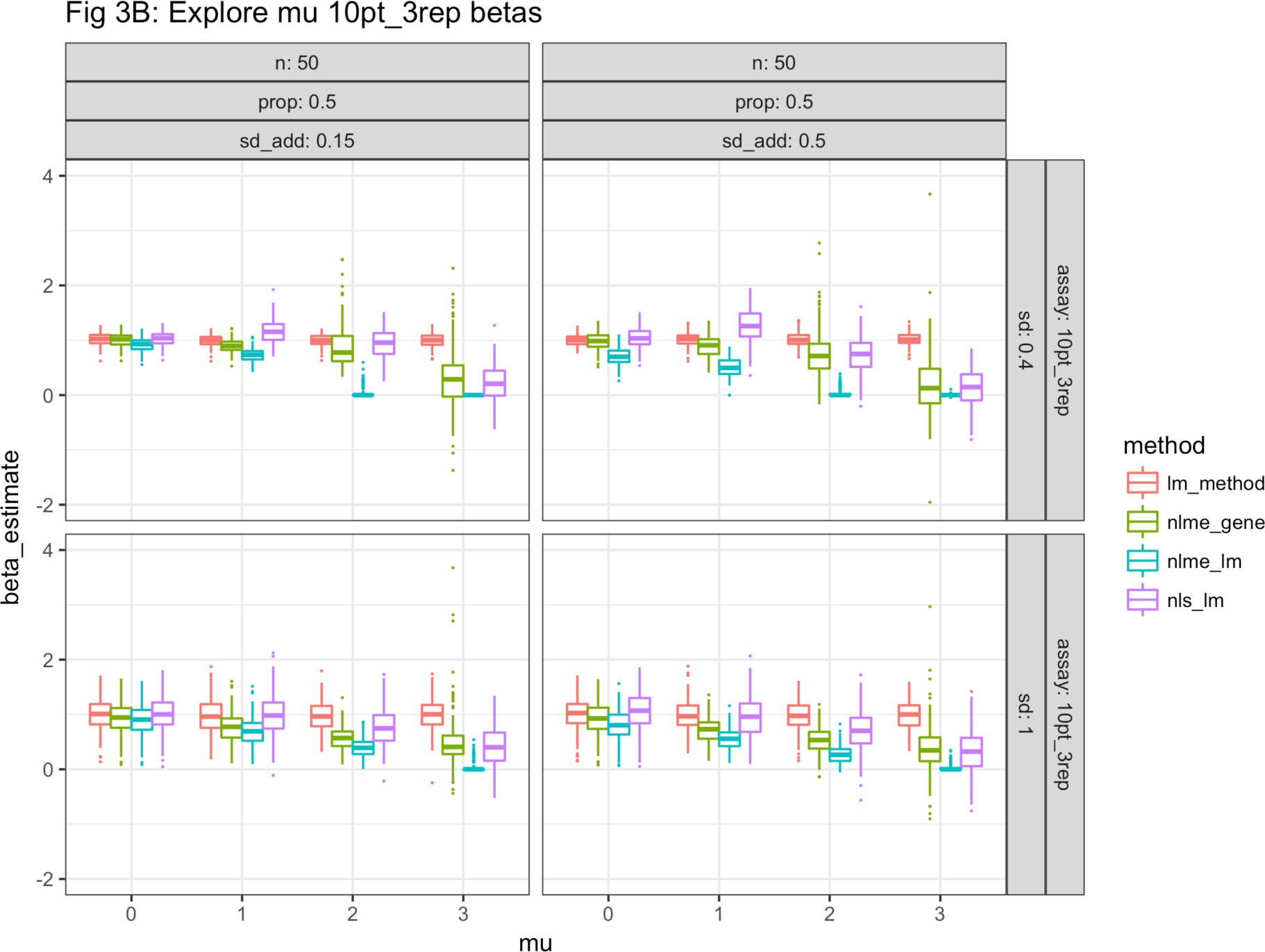
Estimated values of the genetic covariate (beta) for Simulation 2 are plotted on the y-axis in A and B, where A represents results for the 7-point dose response curve and B represents results for the 10-point dose response curve. In C and D the negative log10 of the test-level p-values are plotted on the y-axis for the 7-point and 10-point dose response curves respectively. The x-axis represents the average pIC50. Each panel represents a different set of simulation parameters.

In general, nlme_gene performs the best since it gives the most accurate (or no worse) estimate of the genetic effect size, as well as the most power. In particular, an advantage is seen over the nls_lm method when the average pIC50 is near or above the maximum concentration of the dose response curve, a situation which is often encountered in practice.

### Simulation 3: Exploring error in dose response

Estimated genetic effect sizes and p-values for each simulation are plotted in Figure 4. As in Simulation 2, a systematic shrinkage of effect size estimates is seen using nlme_lm. For low number of cell lines little difference is seen between the methods in terms of power, most likely because there is not enough information to share between cell lines within the mixed effects methodology to improve outcomes. As expected the difference between the baseline lm method and the methods using dose response data increases as noise is added to the dose response data (increasing sd_add), and where there is less information in each dose response curve (7 points single replicate vs 10 points triplicate). In general, however, this simulation shows that the mixed effects approach does no worse than the conventional approach, although there is shrinkage of the effect size estimates in the nlme_lm method, and in some situations performs better.

**Figure 4:**
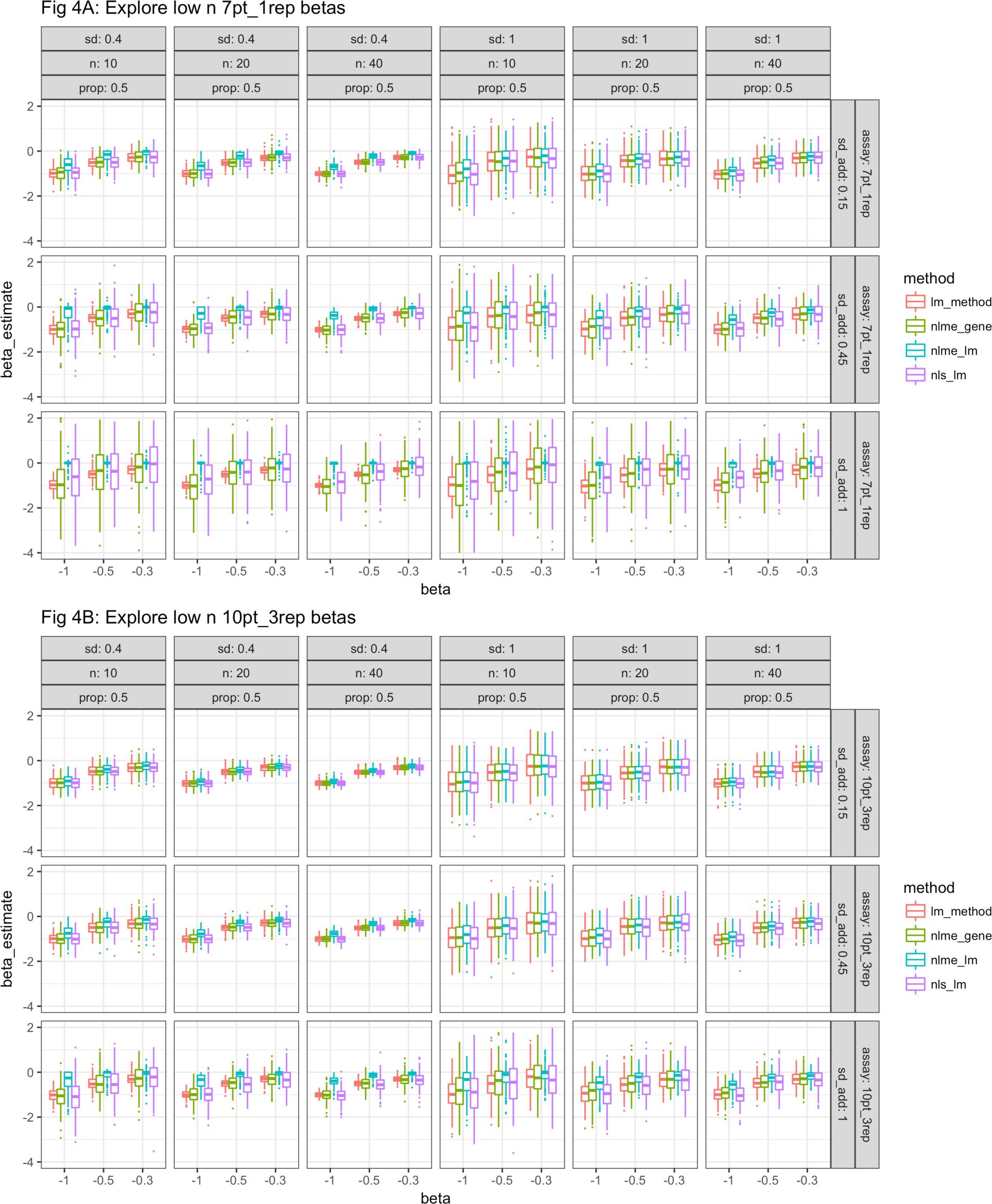
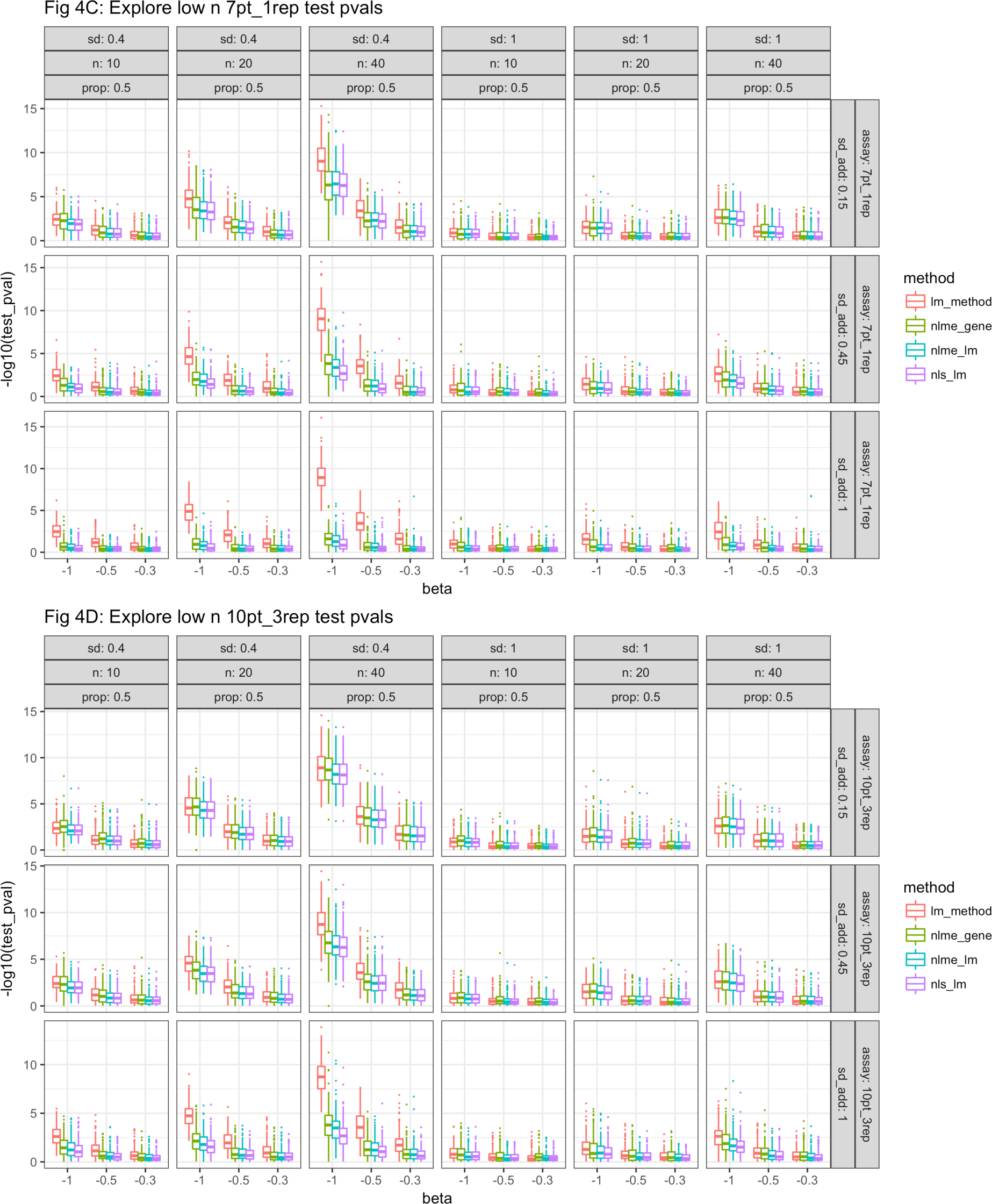
Estimated values of the genetic covariate (beta) for Simulation 3 are plotted on the y-axis in A and B, where A represents results for the 7-point dose response curve and B represents results for the 10-point dose response curve. In C and D the negative log10 of the test-level p-values are plotted on the y-axis for the 7-point and 10-point dose response curves respectively. The x-axis represents the value of the genetic coefficient, beta. Each panel represents a different set of simulation parameters.

### Simulation 4: Exploring experimental design

Estimated genetic effect sizes and test-level p-values for each simulation are plotted in Figure 5 and the results of a power calculation are plotted in Figure 6. As seen previously, there is a shrinkage effect in the estimates of the effect size using nlme_lm. As expected, beta estimates are more precise, and p-values are smaller when more cell lines are simulated and when the effect size is bigger. With the 10-point triplicate dose response curves, the methods perform equivalently well but with the 7-point single replicate dose response curve differences are observed. For instance when the effect size is smaller, and the additive noise is increased, the nlme methods are more powerful than the nls_lm method. This is manifested by the separation of the methods in the power calculation (Figure 6) and the differences in p-values in Figure 5. The power calculation also shows that in general the best way to increase power is to examine more cell lines.

**Figure 5:**
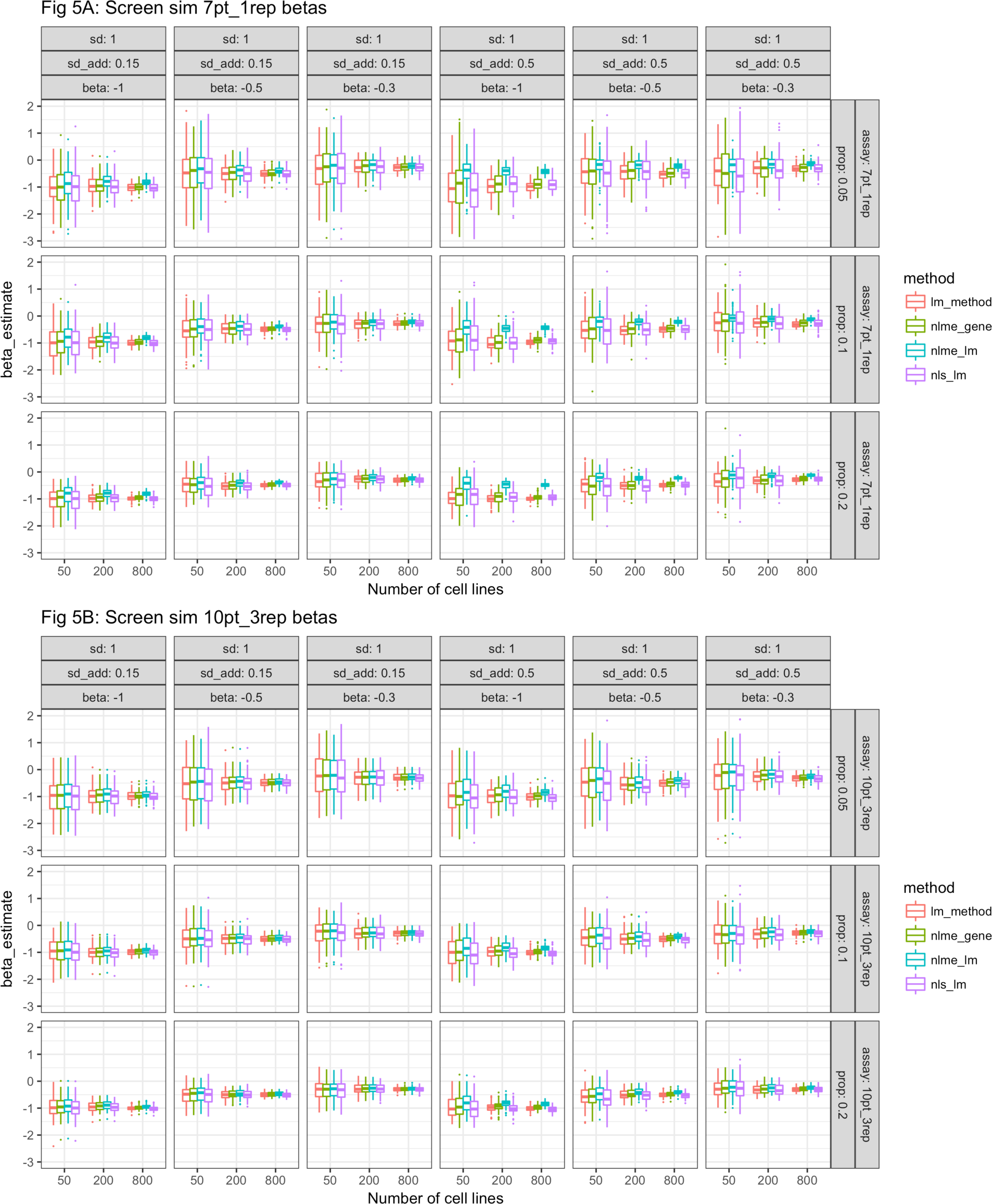
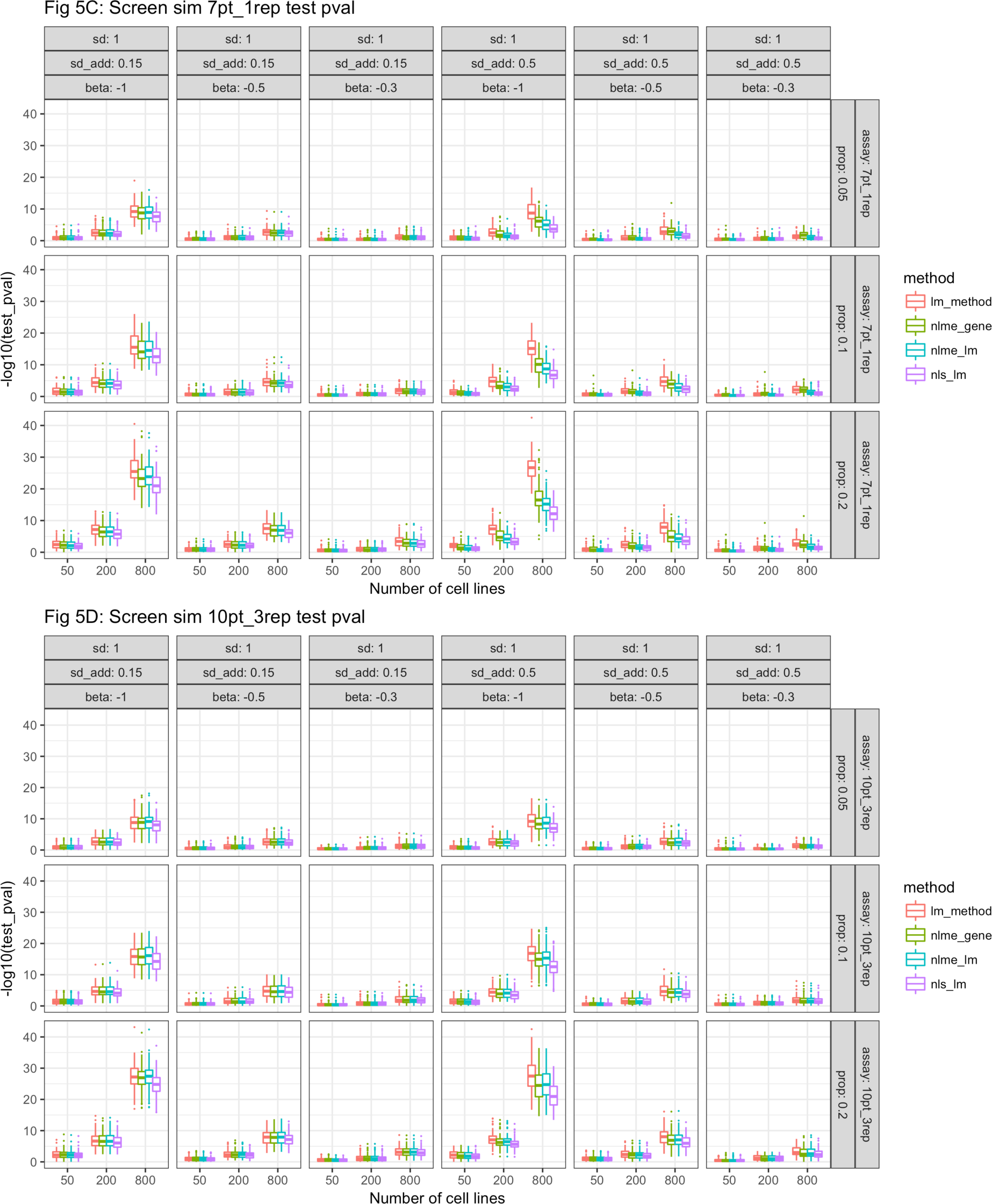
Estimated values of the genetic covariate (beta) for Simulation 4 are plotted on the y-axis in A and B, where A represents results for the 7-point dose response curve and B represents results for the 10-point dose response curve. In C and D the negative log10 of the test-level p-values are plotted on the y-axis for the 7-point and 10-point dose response curves respectively. The x-axis represents the number of cell lines simulated. Each panel represents a different set of simulation parameters.

**Figure 6:**
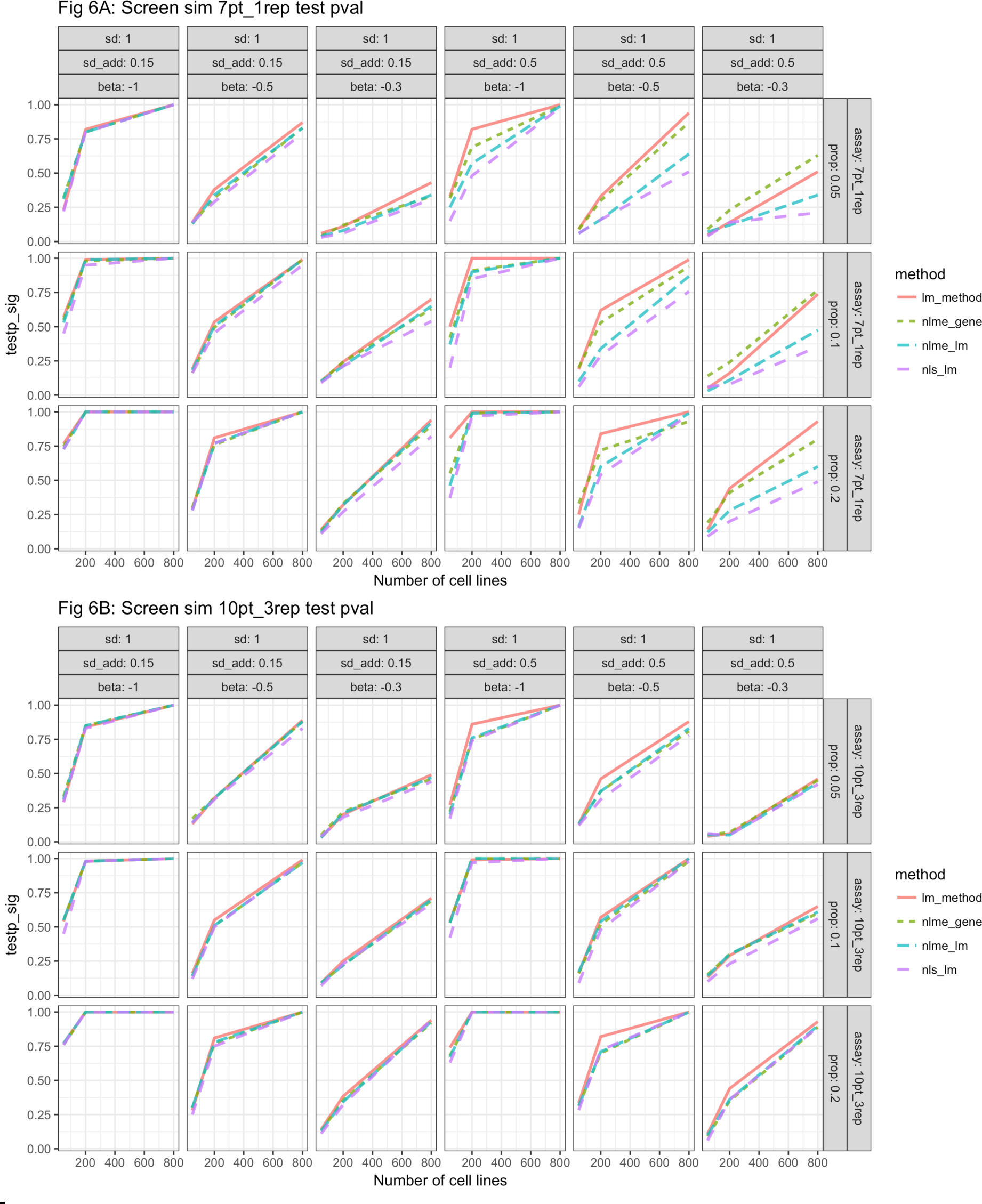
Power calculations for simulation 4. The y-axis represents the proportion of times that the test p-value fell below an arbitrary cut off of 0.05 whereas the x-axis represents the number of cell lines simulated. A represents results for the 7-point dose response curve and B represents results for the 10-point dose response curve. Each panel represents a different set of simulation parameters.

## Discussion

Correlating genomic features to drug sensitivity measures such as IC50 values across a panel of cancer cell-lines is a core activity carried out by pharmacogenomics researchers within both academic and industrial settings. There is a lack of literature within this field on the design of dose-response experiments but an interest in exploring new metrics and analytical methods for such experiments (16,18,20,21). The aim of this study was to show how simulation studies can be used to aid in the design of dose-response experiments and assist in the choice of analysis method to be applied.

Simulation studies are encouraged by the statistical community to assist with understanding the data generation process and the limitations of the planned analysis methods used to analyse real data (cite the statistical rethinking book). Here we set-up a simulation protocol to generate noisy dose-response data for a population of cell lines whose sensitivity to drug is dependent on mutation status. The simulation protocols considered here explored varying degrees of the following: i) number of doses and noise in the dose-response curves; ii) number of technical and biological replicates; iii) proportion and effect size of drug sensitive cell lines within a cell population; and v) population mean and variance of IC50 values.

We considered 3 different analysis methods with increasing technical complexity. The first and simplest approach, defined as nls_lm, involves estimating an IC50 value for each cell line one at a time and then assessing whether the pIC50 (pIC50 = log_10_(IC50)) values correlate to cell-line mutation status using ANOVA. The second, defined as nlme_lm, is similar to the first except that IC50 values are estimated using the non-linear mixed effects method i.e. pooling all dose-response data together to first estimate the population IC50 mean and variance values before estimating each individual cell lines IC50 value (cite Vis et al.). Finally, the third and most technically complex method, defined as nlme_gene, involves exploring the correlation between IC50 values and cell-line mutation status by including mutation status as a covariate within the mixed-effects framework.

From the simulation studies conducted here we found that only under certain conditions did the most complex method, nlme_gene, give us an improvement in statistical power over the simplest method, nls_lm. These cases were when there were fewer data points in the dose response curve and the numbers of cell-lines were high. We found that in situations when the numbers of cell-lines were low, regardless of the number of data points, the simplest approach had just as low power as the most complex approach. We also saw a shrinkage of the genetic effect size in the nlme_lm approach, a phenomenon that has been described previously in the context of pharmacokinetic studies (22). Overall, the nlme_gene method was shown to have increased power when looking across all designs without exhibiting the shrinkage effect. This result makes intuitive sense since the complex method, nlme_gene, uses more information to estimate measurement uncertainty than the simple method, nls_lm.

The simulation methodology and software described here will enable scientists within the pharmacogenomics field to assess the power of different experimental designs and analysis methods. This should enable the community to design better and more cost-effective experiments which will hopefully improve the outcome of analyses.

The main limitations of this study are as follows. First we only considered variation in the IC50 value across cell-lines which may not always be the case since others have shown that both the steepness and the maximum cell death within a dose-response curve can vary across cell-lines under a single drug (21). Second, we have only considered genotype as a discrete covariate (i.e. mutation status) and thus have not considered continuous genomic covariates such as gene expression. Thirdly, we used fixed values of noise for all cell lines which is not likely to be the case as the smoothness of the dose-response curve could be cell line specific. Finally, we have not considered that there could be a correlation structure in the IC50 values across the cell line panel i.e. cell-line sensitivity can vary by tissue (19). All of these limitations can be handled by increasing the complexity of the simulation protocol. However in doing so makes the analysis of the results more complex too.

## Conclusion

In summary the analysis and methodology highlight the value of conducting simulation studies within the field of pharmacogenomics. Within the specific case study relating genotype to cell-line response data we found that more complex methods using the mixed-effects model framework, that are not currently routinely used, can increase the statistical power over the current methods. We hope this encourages the pharmacogenomics community to conduct more simulation studies but also build on the work described here.

## Acknowledgements

- Louis Aslett (Durham) for RStudio AMI’s

